# An open-access volume electron microscopy atlas of whole cells and tissues

**DOI:** 10.1101/2020.11.13.382457

**Authors:** C. Shan Xu, Song Pang, Gleb Shtengel, Andreas Müller, Alex T. Ritter, Huxley K. Hoffman, Shin-ya Takemura, Zhiyuan Lu, H. Amalia Pasolli, Nirmala Iyer, Jeeyun Chung, Davis Bennett, Aubrey V. Weigel, Melanie Freeman, Schuyler B. van Engelenburg, Tobias C. Walther, Robert V. Farese, Jennifer Lippincott-Schwartz, Ira Mellman, Michele Solimena, Harald F. Hess

## Abstract

Understanding cellular architecture is essential for understanding biology. Electron microscopy (EM) uniquely visualizes cellular structures with nanometer resolution. However, traditional methods, such as thin-section EM or EM tomography, have limitations inasmuch as they only visualize a single slice or a relatively small volume of the cell, respectively. Focused Ion Beam-Scanning Electron Microscopy (FIB-SEM) demonstrated the ability to image cellular samples at 4-nm isotropic voxels with rather limited imageable volume. Here, we present 3D EM images of whole cells and tissues with two orders of magnitude increases in imageable volume at 4-nm voxels. Such data with a combined fine resolution scale and large sample size do not currently exist, and are enabled by the advances in higher precision and stability of FIB milling, together with enhanced signal detection and faster SEM scanning. More importantly, we have generated a volume EM atlas encompassing ten diverse datasets of whole cells and tissues, from cancer cells to immune cells, and from mouse pancreatic islets to *Drosophila* neural tissues. These open-access data (via <u>OpenOrganelle)</u> represent a foundation to nucleate a new field of high-resolution whole-cell volume EM and subsequent analyses, and invite biologists to explore this new paradigm and pose fundamentally new questions.

---

Individual cells and tissues can be described by a 3D hierarchy of ultrastructural details, from single protein molecules to organelles within complete cellular architecture^1^. Transmission electron microscopy (TEM) images from ∼30–300 nm diamond-knife cut sections of metal stained samples provide convenient high-resolution visualization on a small fraction of the whole cell, which have profoundly shaped our understanding of organelles^2^. Series of sections can be digitally stitched together to form a 3D volume, but the 30 nm or larger sampling interval sacrifices structural detail. Tilt tomography achieves higher 3D resolution on ∼200-nm-thick sections. Alternatively, diamond-knife cut surfaces of a block face sequentially imaged with scanning electron microscopy (SEM)^3^ was a pioneering step to access larger volumes, but its z resolution is limited to 25 nm due to electron radiation damage from beam scanning that prevents consistent cutting of thinner slices^4^ in subsequent rounds. This z-resolution constraint can be overcome using FIB-SEM that offers high isotropic resolution. However, its deficiencies in imaging speed and duration (several days) cap the maximum imageable volume. Our prior work (Fig. 1a) reported the transformation of a conventional FIB-SEM lacking long-term reliability into an enhanced platform capable of months to years of continuous imaging at 8-nm isotropic voxels without defects in the final image stack^5, 6, 7, 8^, but we struggled to obtain volumes of ∼500 µm^3^ at 4-nm voxels due to slow SEM scan using backscattered electron detection via a sample biasing scheme^5, 9^, and FIB milling instability induced by higher electron radiation energy density (∼400 keV/nm^3^). Thus, higher-resolution isotropic 3D whole cell data remained mostly out of reach.

**Fig. 1.**
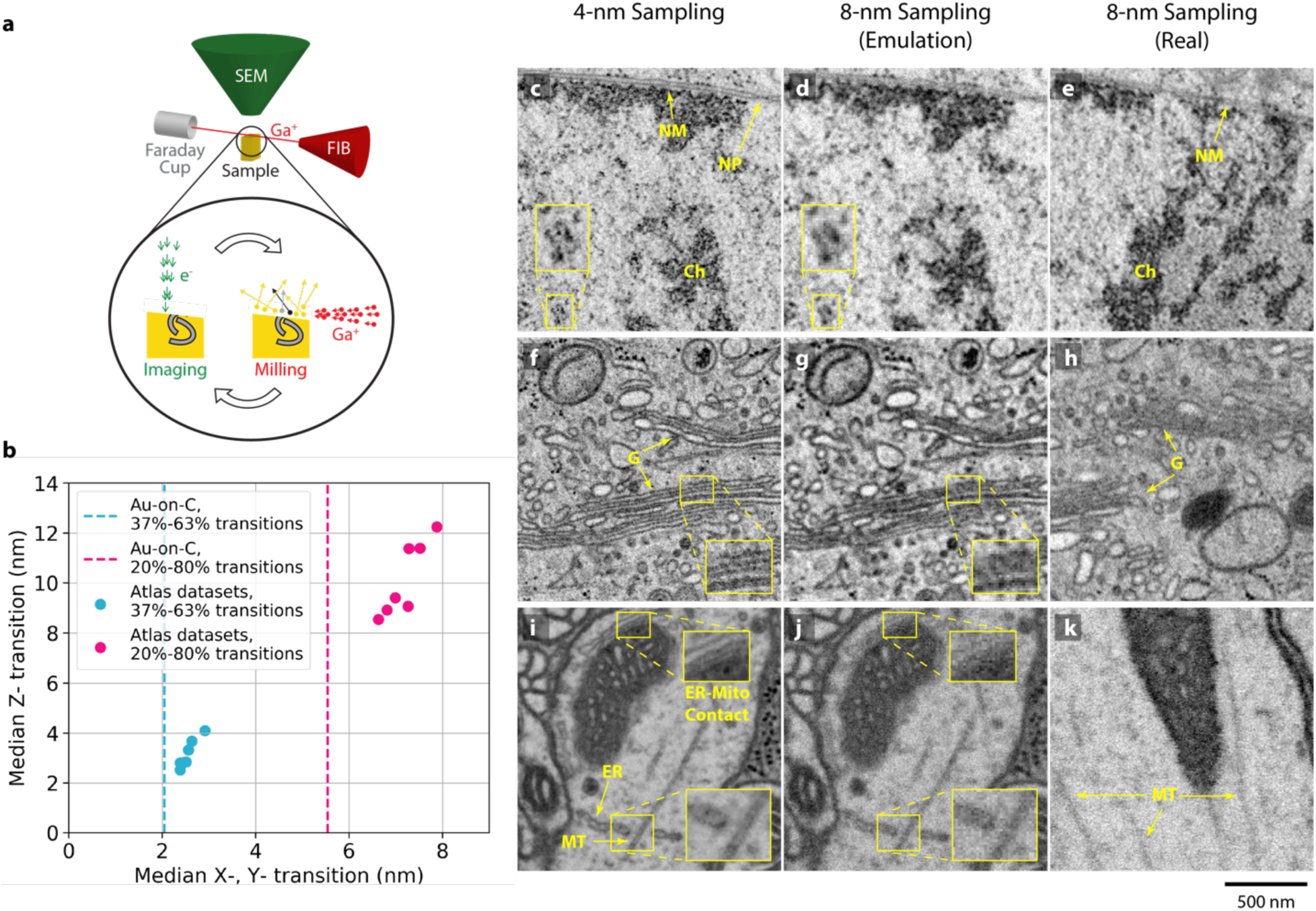
Enhanced FIB-SEM configuration, operation, and resolution. **a**, Sketch of FIB milling and SEM imaging. The two operations iterate alternately to generate 3D whole cell image stacks, **b**, Resolution characterization using transitions at the edges of gold nanoparticles on a carbon substrate and ribosomes in cultured cells. The vertical dash lines indicate the edge transitions of gold nanoparticles, the round dots indicate the edge transitions of ribosomes along x-y and z axes. Each dot represents the average value from more than 3000 ribosomes in one of the seven cultured cell samples. **c–k**, Comparison of lower current/higher resolution 4-nm sampling and 8-nm sampling. Left column shows images acquired at 4-nm sampling that matches the resolutions shown in **b**. The middle column shows the emulated 8-nm images (details in Methods Section), while the right column shows the real 8-nm sampling images. **c**, Nucleus of the interphase HeLa cell from Fig. 2d shows fine details of chromatin (Ch), nuclear membrane (NM), and nuclear pore (NP). Nucleosomes are less resolved in **d,** the emulated 8-nm sampling image, and in **e**, the real 8-nm sampling image of an interphase HeLa cell. **f**, Golgi (G) cisternae of the interphase HeLa cell from Fig. 2c are better resolved at 4-nm sampling compared to the emulated 8-nm sampling image shown in **g** and the interphase HeLa cell imaged at 8-nm sampling shown in **h**. **i**, *Drosophila* brain sample from Supplementary Video 1 shows well-resolved hollow core of a microtubule (MT) and the close contract between endoplasmic reticulum (ER) and mitochondria (Mito), which are not distinguishable in **j**, the emulated 8-nm sampling image. **k**, The real 8-nm sampling image of blurry microtubules of a *Drosophila* brain sample with higher shot noise and less resolved microtubules compared to those shown in **j**. Scale bar, 500 nm in all images.

Here we report imageable volumes greater than 100,000 µm^3^ at 4-nm voxels. This newly accessible paradigm is illustrated in Extended Data Fig. 1. The two orders of magnitude improvement in volumes compared to our prior work^5^ primarily come from: 1) higher precision and stability of FIB milling control to extend reliable long-term acquisition to 4-nm voxels, and enhanced SEM signal detection using secondary electrons with higher beam current to achieve faster imaging, hence reducing electron radiation energy density (from ∼400 to ∼80 keV/nm^3^) that further improves FIB milling control. To overcome milling instability, we integrated a more stable FIB column (Capella from Zeiss), and tightened its FIB emission current control band to obtain a consistent milling beam profile. We also re-configured the closed-loop control^5^ to reduce the milling variation between FIB reheat cycles. Such FIB milling optimization allowed for faster SEM imaging^5^ using secondary electrons with 5x lower electron radiation energy density, which subsequentially further mitigated FIB milling artifacts and instability. Additionally, we investigated a multi-dimensional operation space to optimize image contrast and isotropic resolution. We found that a beam current of 200 to 300 pA and a landing energy of 700 to 900 eV are optimal which further accelerate the imaging rate. Such positive synergy between faster SEM scanning and robust FIB milling has extended reliable imaging acquisition at 4-nm isotropic voxels from less than a week to months. The detailed instrumentation improvements are described in the Methods Section. To validate the choice of sampling voxel size, we characterized the resolution in x-y and z by analyzing the transitions at the edges of gold nanoparticles on a carbon substrate and ribosomes in cultured cells. The ribosomes in biological specimens serve as a convenient *in situ* resolution standard. Fig. 1b shows near-isotropic resolution matching of the 4-nm voxel size with the average transitions from 37% to 63% being 2.5 nm in x-y and 3.1 nm in z, respectively. The x-y transitions determined from ribosome edges are about 30% larger than those for the gold nanoparticles (vertical lines in Fig. 1b). The procedure for estimating the resolution with examples is presented in the Methods Section. Furthermore, an improved sample staining protocol yielding higher contrast leads to additional 5x–10x scanning rate improvement, thereby 200x larger volume is demonstrated on *Drosophila* brain samples (Supplementary Video 1). The maximum volume can be seamlessly extended with longer imaging acquisition.

Enabled by these advances, we here present a 3D atlas of whole cells and tissues at 4-nm voxels. The initial datasets (Table 1) consist of ten wild-type biological specimens from cancer cells to immune cells, and from mouse pancreatic islets to *Drosophila* neural tissues, each requiring weeks of uninterrupted imaging. The resulting data demonstrate the finest possible isotropic resolution for whole cell volumes. To highlight some features that are only visible at the higher resolution, a comparison between 4-nm versus 8-nm voxel sampling is illustrated in Fig. 1c–k. In particular, the nucleosomes inside its nucleus showing finer details are on the verge of being individually resolved at 4 nm (Fig. 1c); the detailed Golgi cisternae (Fig. 1f), the close contacts of ER-mitochondria, and the hollow core of microtubules (Fig. 1i) are only resolvable at 4 nm. Furthermore, Supplementary Video 1 visualizes the value of finer resolution with 200x larger volume to better illustrate the newly accessible combined larger size-higher resolution paradigm. By making these data openly available, we invite biologists to contemplate analyzing such comprehensive whole cell details, and hope to inspire new questions that harness complete rather than partial cell data to unveil new insights.

**Table 1.**
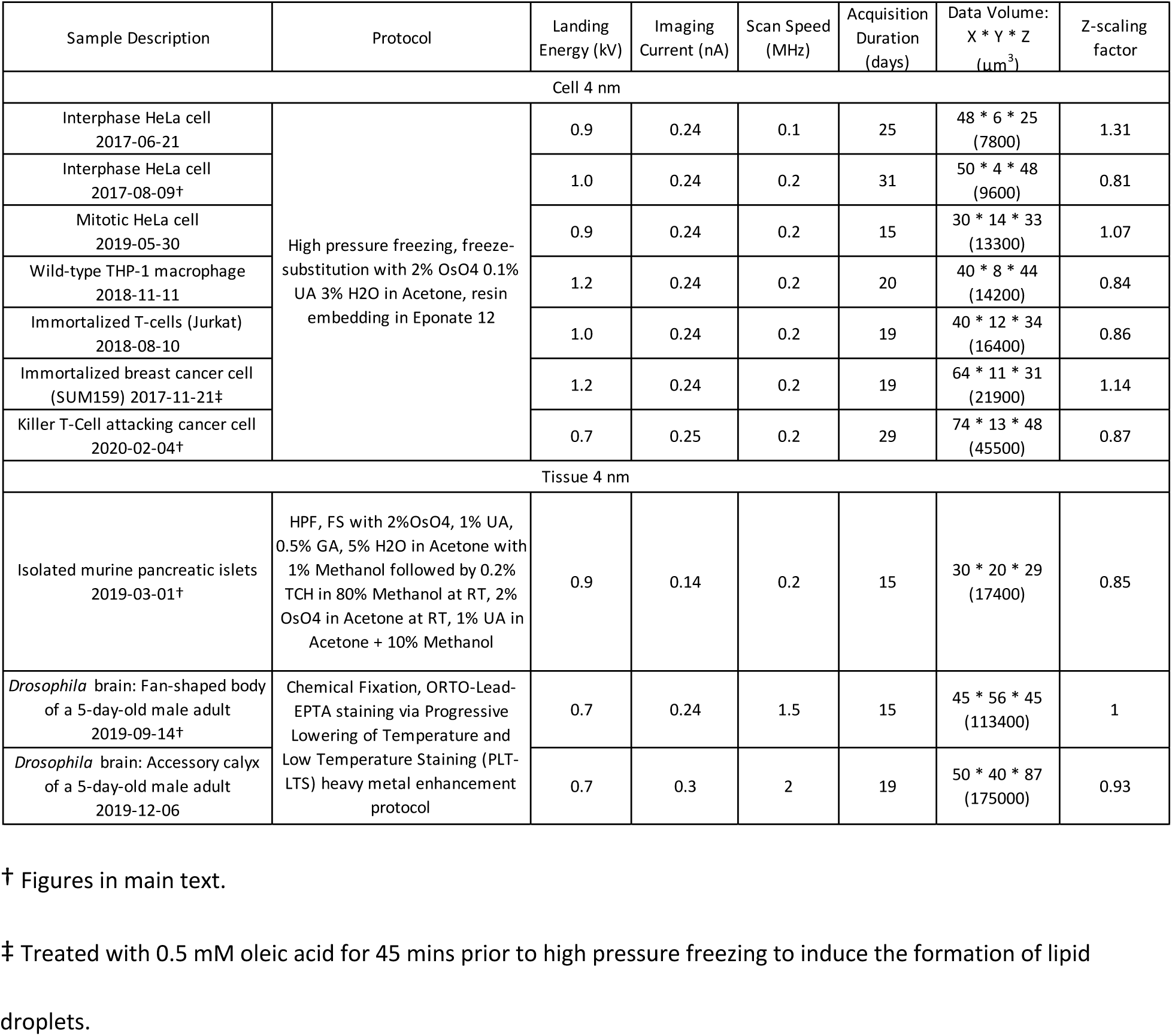
Descriptions of exemplary datasets with selected imaging conditions

**HeLa cells**, human cervical cancer cells, are easily cultured and widely used in cell biology laboratories as a basic model to test diverse hypotheses. Fig. 2 presents a typical 3D dataset of an entire HeLa cell. The top panel provides an overview with manually segmented mitochondrial network in green (Fig. 2a). The bottom row panels show 2D cross-sections of other cellular organelles, such as the centrosome (Fig. 2b), the Golgi apparatus (Fig. 2c), and nuclear envelope (Fig. 2d). No single 2D cross-section allows visualizing all centriole sub-distal appendages, however quick segmentation of the 3D dataset depicts them clearly (red, Fig. 2b). Stereotypical 2D images of the Golgi stacks do not reveal the fenestration details nor long thin tubular extensions that can readily be seen in 3D segmented single Golgi cisterna (magenta, Fig. 2c). Polyribosome chains on the nuclear envelope are mostly hidden in 2D cross-sections, but easily resolved and detailed in 3D (yellow, Fig. 2d). Evidently, having isotropic 3D images of such cells in their entirety can provide an accurate reference to which perturbations in growth, genetic, environment, etc. can be compared.

**Fig. 2.**
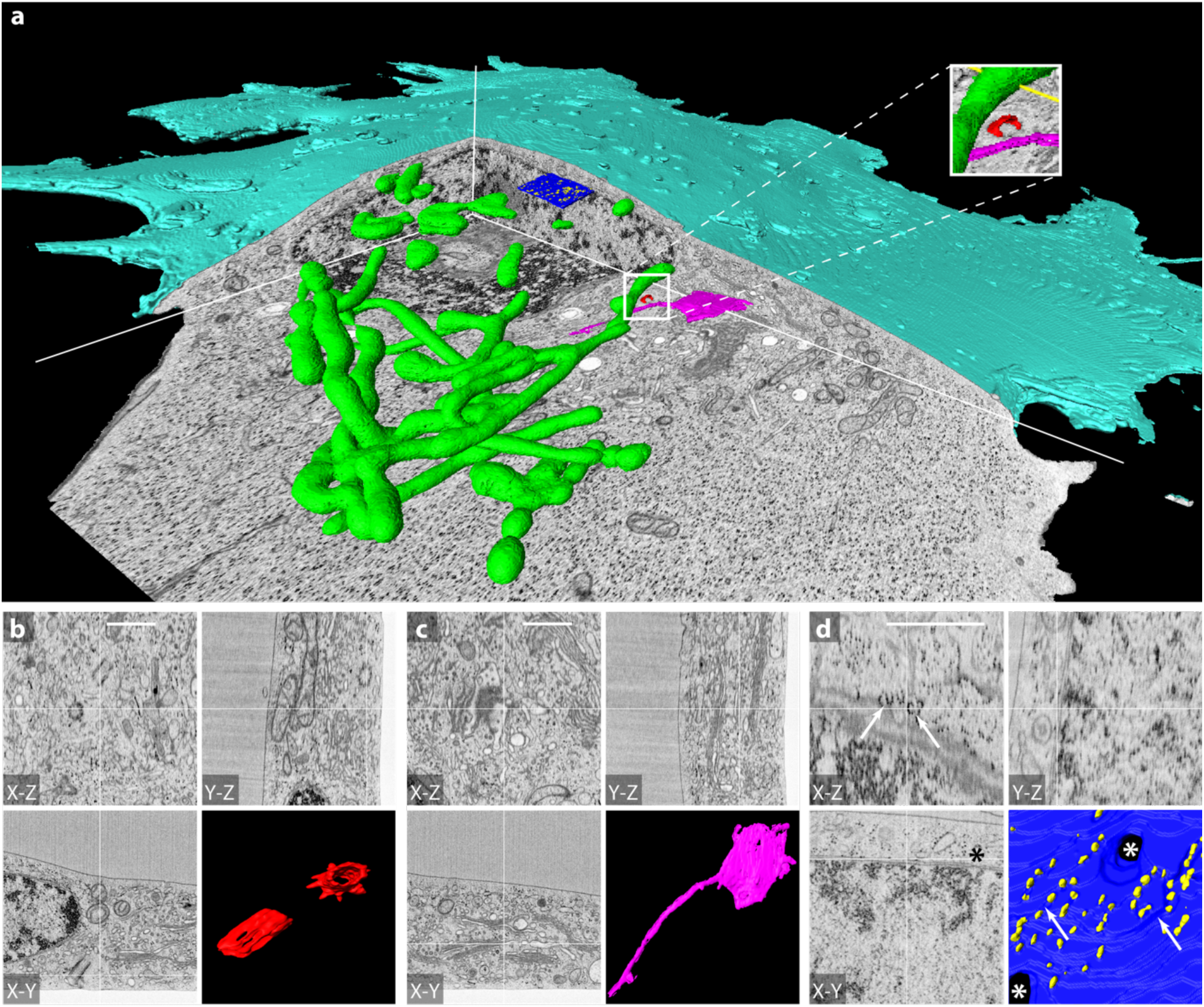
Interphase HeLa cell. **a**, FIB-SEM overview with cutaway, and manually segmented interior features (mitochondria, green; centrosomes, red; one cistern of a Golgi stack, magenta; a segment of nuclear membrane, blue, polyribosome chains, yellow). Zoomed in cross-sections of three select areas containing: **b**, Centrosomes, **c**, Golgi stack, and **d**, Nucleus, polyribosomes indicated by arrows, and nuclear pores indicated by asterisks. Scale bars are 1 μm.

**Cytotoxic T lymphocytes** (CTLs) are immune cells that have the capacity to identify and destroy virus-infected or cancerous cells. Upon encountering a target, CTLs form an organized intercellular interface called the immunological synapse (IS). Signaling at the IS directs the polarized release of toxic proteins housed in specialized secretory lysosomes called lytic granules (LGs), which results in target cell death. Due to the important role of CTLs in anti-viral and anti-cancer immunity, the IS has been the subject of intense scrutiny. Limited single slice TEM images of CTL:target conjugates have revealed interesting features of the IS, but do not represent the full scope of this dynamic, three-dimensional structure^10^. Fig. 3 presents a first comprehensive 3D dataset of a mature murine CTL (green) engaging an ID8 ovarian cancer cell (cyan). Although there is a minor membrane damage in the area of IS, which most likely occurred during high-pressure freezing or freeze-substitution or resin embedding^11^, the isotropic high-resolution information of FIB-SEM imaging provides a unique and complete map of the complex membrane topology at the interface between T cell and target. Fig. 3b demonstrates the stereotypical CTL “cupping” of the target cell at the IS through ortho-slices. Zoomed in images further illustrate the variety of features across this interface including membrane interdigitation (Extended Data Fig. 2a), flat membrane apposition (Extended Data Fig. 2b), and filopodia of the target cell trapped between the two cells (Extended Data Fig. 2c). Without the ability of high resolution 3D imaging, these filopodia could be mistaken for vesicles trapped between T cell and target. Additionally, Supplementary Video 2 depicts the morphological features in greater details, which further highlights the hallmarks of CTL killing: T-cell cupping the target cancer cell (Fig. 3b), polarization of the centrosome toward the target cell^12, 13^ (Fig. 3c), and the lytic granules with diverse ultrastructures aggregating near the synapse (Fig. 3d). Such high resolution whole cell FIB-SEM dataset allows for the label-free localization of all organelles in the T cell including those responsible for target killing and cytokine secretion.

**Fig. 3.**
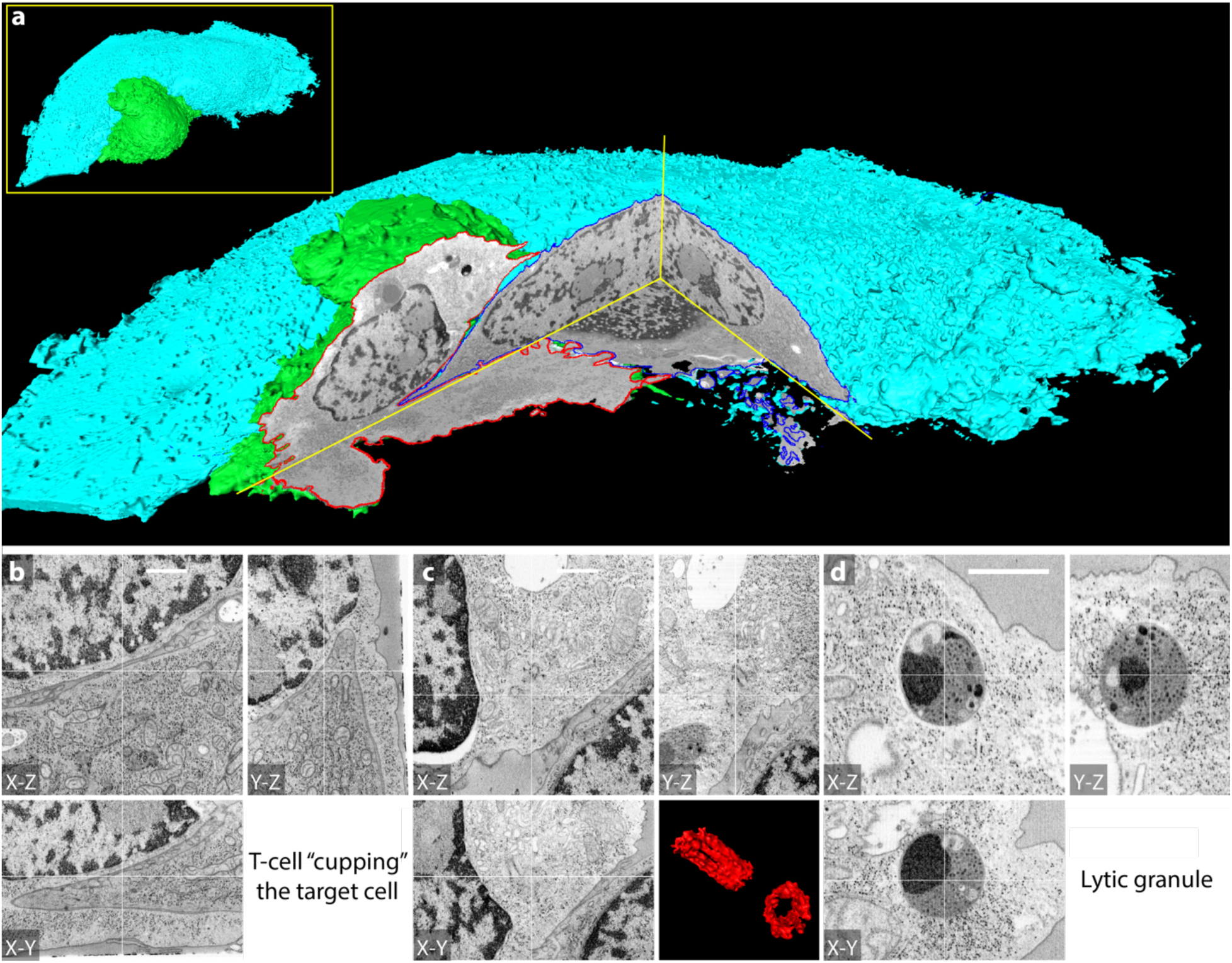
Murine CTL engaging an ovarian cancer cell. **a,** FIB-SEM overview with cutaway, and manually segmented membranes of CTL (green surface and red contour) and cancer cell (cyan surface and blue contour). Zoomed in cross-sections highlight the signatures of a productive immunological synapse: **b**, CTL cell “cupping” the target cancer cell, **c,** Polarized centrosome, and **d**, Lytic granule showing unique ultrastructures. Scale bars are 1 μm.

**Pancreatic islets** are micro-organs consisting mainly of beta, alpha, delta, polypeptide cells and endothelial cells. Among them beta cells are in the majority and secrete insulin stored in secretory granules (SGs) to maintain blood glucose homeostasis. Few beta cells have been reconstructed by serial section electron tomography at medium resolution^14^ to allow assessment of subcellular features such as insulin SG number and shape of mitochondria, but lacking the resolution for imaging of e.g., ER and microtubules. Large-scale high-resolution FIB-SEM enables the analysis of ultrastructural differences between beta cells within an islet as well as features that require higher resolution such as ribosomes and the cytoskeleton. Fig. 4a displays several complete beta cells that are entirely included within the 30 x 20 x 25 µm^3^ volume. The large volume data allow for studying the interaction of beta cells, e.g., full primary cilia and their connections to neighboring cells and the intermingling of microvilli (Fig. 4b, 4c). At 4-nm voxel sampling, we observed ultrastructural differences between insulin SGs (Fig. 4d). For the first time, we report close contacts of ER and insulin SGs in 3D (Fig. 4e). Ultimately, such high resolution allows the sharp definition of microtubules, single ribosomes, membrane vesicles, and Golgi cisternae for quantitative measurements (Fig. 4f). The seamless imaging results in precisely aligned stacks, further facilitating automated 3D segmentation. Specifically, emerging from these FIB-SEM datasets is a comprehensive 3D representation of microtubule networks and how they interact with other organelles^15^. We could show that beta cell microtubules are arranged into non-radial networks which for the most part do not originate either from centrioles or endomembranes^15^. Moreover, glucose stimulation changes the number and length of microtubule polymers without changing their overall density^15^. Finally, we found insulin SGs to be enriched near the plasma membrane in association with microtubules, independent of the extracellular glucose levels^15^.

**Fig. 4.**
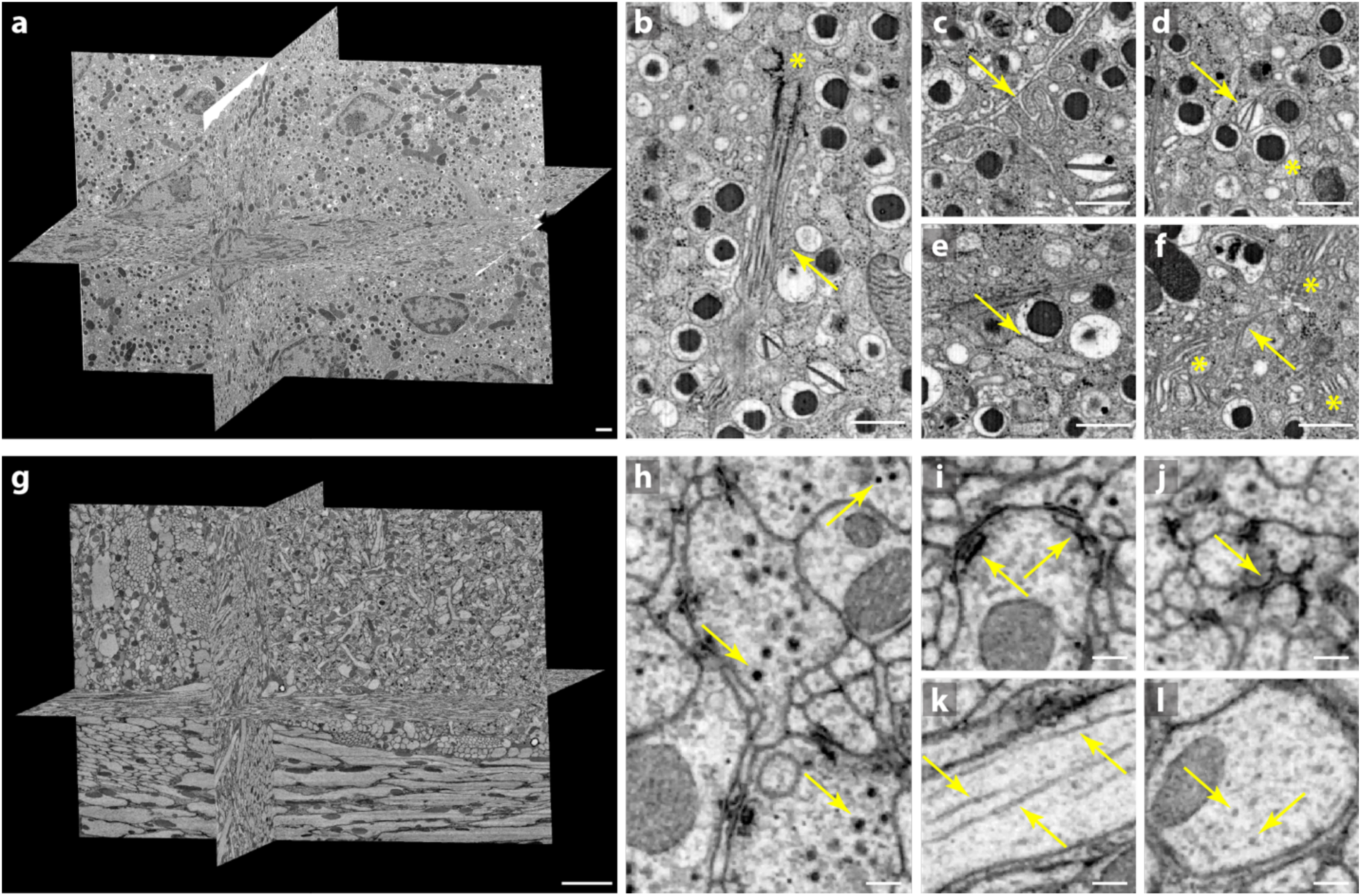
Tissue sample datasets. **a**, Pancreatic islets treated with 16.7 mM glucose containing several complete beta cells with detailed structures shown in **b**, A primary cilium with axoneme (arrow) and centrioles (asterisk), **c**, Intermingled microvilli (arrow), **d**, Ultrastructural diversity among insulin SGs containing rod-shaped (arrow) or spherical crystals (asterisk), **e**, Contacts of ER and insulin SGs (arrow), **f**, Microtubules (arrow), Golgi apparatus (asterisks) and ribosomes. **g**, *Drosophila* fan-shaped body with detailed structures shown in **h**, Dense-core vesicles of different sizes (arrows), **i**, Multiple synaptic sites viewed from the side (arrows), **j**, A presynaptic T-bar viewed from the top (arrow), **k**, A longitudinal-section view of microtubules (arrows), **l**, A cross-section view of microtubule arrays (arrows). Scale bar, 1 µm in **a**, 500 nm in **b**–**f**, 5 µm in **g**, 200 nm in **h**–**l**.

The **central complex in insect brains** is a highly conserved brain region that controls various behaviors. Fig. 4g shows higher-resolution images acquired from the fan-shaped body of the central complex of *Drosophila*. To understand brain functions, it is imperative to identify the synaptic levels of neuron connectivity over large volumes through key synaptic elements such as presynaptic T-bars and postsynaptic densities (PSDs) (Fig. 4i, 4j). To investigate if the central complex contains characteristic synaptic motifs, such as polyadic synapse, rosette synapse, or any new synaptic motif, validating the synaptic structures with high-resolution data is crucial. However, a trade-off between volume and resolution often limits capabilities in connectomics studies, where large-scale neuronal tracing and fine synaptic structure identification are equally important. High resolution large volume FIB-SEM imaging, shown here, bridges these capabilities and therefore provides significant possibilities. Such datasets allow reconstruction of not only the large neuronal objects and glial cells, but also smaller intracellular components such as synaptic vesicles, dense-core vesicles (Fig. 4h), and microtubules (Fig. 4k, 4l). Knowing the types of synaptic vesicles could add important insights to understand the properties of synaptic connections. As an example, the terminals of dopaminergic neurons in the fly’s olfactory center were shown to contain both clear and dense-core vesicles^16^, in congruence with having more than one neurotransmitter^17^.

A considerable investment in technology development, equipment resources, and imaging duration is required to produce large volume high resolution datasets, but the resulting wealth of such data is unprecedented and comprehensive, covering whole cells with all their organelles and proteins that can be highlighted with heavy metal stains. Here, we open access to this volume EM atlas through <u>OpenOrganelle</u> (https://openorganelle.janelia.org)^18^. With access to these datasets and future ones, biologists can assess whether whole cell structure statistics and architecture characterization of their interest are visible and possible to measure for evaluating the merits of further exploration. Additionally, the data can be used as a reference ultrastructural atlas, providing insights into both known and previously unappreciated structural details of intracellular organization. Each dataset can be downloaded to mine, while pre-selected views of interest and cross-sections can be browsed on-line with *neuroglancer*, a built-in visualization tool. More importantly, these data enable a holistic way of looking into the cellular world and stand as a foundation of segmentation and analysis efforts that could be pursued by the scientific community.

In summary, we achieved large imaging volumes over 100,000 µm^3^ at the finest possible isotropic resolution. Thereby, entire mammalian cells and multicellular parts of tissues were imaged at 4-nm isotropic voxels to reveal a more holistic 3D cellular architecture. Ten diverse datasets of reference wild-type examples are now made accessible through <u>OpenOrganelle</u>^18^. This allows the study of complex cell morphology, complete interfaces involved in cell-cell interactions, and the shapes, volumes, distributions, and relationships of intracellular organelles. Particularly, the examination of a wild-type cell can encourage further experiments to compare features against perturbations of disease, mutations, development, or environment at the whole cell level. These datasets and targeted feature analyses not only could lead to discoveries of unanticipated new features, but also should spur developments of general tools to break the bottleneck of 3D data mining. Automation of the time-consuming process of segmentation on the whole cell scale is a crucial next step. Such an approach is presented in the associated paper^18^. We believe that open access to rich 3D FIB-SEM datasets will provide important insights, guidance, and inspiration for the research community to further explore and address a new broad range of biological questions.

## Methods

The initial datasets of seven common wild-type cultured cells (from cancer cells to immune cells) and three tissue samples (from mouse pancreatic islets to *Drosophila* neural tissues) are tabulated in Table 1. Detailed procedures are described below.

### Cultured cell sample preparation

Cultured cells were cryo-fixed by high pressure freezing (HPF), which vitrified a sample on a millisecond time scale to best preserve any dynamic structural details. Although HPF sometimes can cause segregation artifacts (i.e. freezing damage), it minimizes any possibility of artifacts that might be of concern with chemical fixation. HPF remains the only viable option for non-chemical fixation of samples more than a few hundred nanometers thick. The subsequent freeze substitution labeled biomolecules with heavy metals to provide contrast for EM.

#### 1. HeLa cells

HeLa cells (CCL-2) were purchased from American Type Culture Collection (ATCC). HeLa cells were maintained in EMEM medium (ATCC, 30-2003) supplemented with 10% FBS (Corning, 35-011-CV) and 1X penicillin-streptomycin solution (Corning, 30-002-CI). Trypsinized HeLa cells were seeded on pre-cleaned and edge-coated sapphire coverslips. Immediately prior to freezing, live cells were inspected to ensure cell morphology and viability. After quality assurance the cells were transferred to a water jacketed CO_2_ incubator (Thermo Fisher Scientific, Midi 40) kept at 37° C, 5% CO_2_, and 100% humidity while awaiting freezing. Each coverslip was removed from the incubator immediately prior to the freezing procedure, and dipped three times in the 25% w/v mixture of Bovine Serum Albumin (BSA) (B4287, Sigma Aldrich)) in the cell medium, which served as a cryo-protectant to prevent ice crystal formation^19^. The samples were then high-pressure frozen between two Aluminum planchettes (Technotrade International, 389 and 479) in Wohlwend HPF Compact 02 High Pressure Freezing Machine (Technotrade International). Once the samples were frozen, they could be stored indefinitely in liquid nitrogen for subsequent freeze-substitution (FS) and resin embedding (RE). FS was performed using automated FS machine (AFS2, Leica Microsystems): coverslips were transferred to cryotubes containing FS media (2% OsO_4_, 0.1% Uranyl Acetate (UA), and 3% water in acetone) under liquid nitrogen, followed by a programmed FS schedule. Resin embedding was performed immediately after FS. Samples were removed from the AFS2 machine, washed 3 times in anhydrous acetone for a total of 10 min and embedded in Eponate 12 which was polymerized for 48 hours at 60°C. Following EPON embedding, the coverslip was separated from the resin block containing the cells by sequential immersion in liquid nitrogen and hot water. EPON was chosen as the initial embedding resin due to less embedding artifacts. However, EPON generates streaking artifact during FIB-SEM imaging^5^. A thin layer (5–10 µm) of Durcupan on the specimen surface facing the FIB beam can effectively mitigate such streaks during FIB-SEM imaging. Therefore, once the coverslip was removed, the exposed surface was immediately re-embedded in Durcupan ACM resin (Sigma Aldrich, set 44610). The detail procedures of coverslip preparation, high pressure freezing, free substitution, and resin embedding are presented in the Supplementary Information.

#### 2. Cytotoxic T cell and T cell/Cancer cell conjugate (CTLs)

To generate CTLs from OT-I mice (Jackson Labs), splenocytes were isolated and stimulated with 10nM OVA257-264 peptide (AnaSpec, Fremont, CA, USA) in 10% RPMI (RPMI 1640 plus 10% fetal bovine serum (Fisher), 2mM L-glutamine, 50U/mL penicillin/streptomycin, and 50μM β-mercaptoethanol. Following 3 days of stimulation, cells were resuspended in complete media plus 10 IU/mL recombinant human IL-2 (rHIL-2, Roche), and seeded in fresh media at 0.5×10^6^ cells/mL every 48 hours. ID8 cancer cell line (ATCC) was grown in RPMI media (Gibco) with 10% FBS (Gibco).

In-vitro activated CTLs from Ova-transgenic (OT-I) mice were combined with adherent ID8 murine ovarian cancer cells. To facilitate recognition of cancer cells by the OTI-I CTLs, ID8 cells suspended in complete medium (RPMI with 10% FBS, Gibco) were incubated with OVA_257-264_ SIINFEKL peptide for 1 hour at 37°C. The cells were washed three times with complete medium, and 10^5^ cells were added to each well of a 24-well plate. Each well contained a single sapphire coverslip coated with human Fibronectin (Corning). The ID8 cells were allowed to settle and adhere for 2 hours in an incubator at 37°C/5% CO_2_. At this time, the media in the well was replaced with 250uL phenol-red free RPMI (Gibco). Care was taken to ensure that media and cells were stored in 37°C/5% CO_2_ for all incubation periods. Immediately after changing the media, 10^5^ CTLs in 50uL complete medium were added to the well. The CTLs were allowed 7 minutes to find their targets and secrete lytic granules before fixation through high pressure freezing. The relatively short incubation time favors capture of CTL:target conjugates in an early-stage of interaction.

The remaining cryofixation, FS, and resin embedding procedures were identical to those described for HeLa cells. The detail procedures are presented in the Supplementary Information.

### Tissue sample preparation

#### 1. Mouse pancreatic islets

Pancreatic islets of 9-week-old C57BL/6 mice were isolated as previously described^20^. They were cultured overnight in standard culture media (RPMI 1640 (Gibco) with 10% FBS, 20 mM HEPES, 100 U/ml each penicillin and streptomycin) containing 5.5 mM glucose. Prior to high pressure freezing the islets were subjected to 1 hr incubation in Krebs-Ringer buffer containing either 3.3 mM or 16.7 mM glucose.

Islets were frozen with a Leica EM ICE high pressure freezer (Leica Microsystems, Germany) and kept in liquid nitrogen until freeze substitution. Although freezing damage was observed in couple nuclei, the preservation of cytoskeletal elements benefits from this fixation. High pressure frozen islets were substituted as previously published^21^ or according to a novel protocol: first the samples were substituted in a cocktail containing 2% OsO_4_, 1% UA, 0.5% glutaraldehyde, 5% H2O in Acetone with 1% Methanol at −90°C for 24 hours. The temperature was raised to 0°C over 15 hours followed by 4 washes with 100% acetone for 15 min each and an increase in temperature to +22°C. Afterwards the samples were incubated in 0.2% thiocarbohydrazide in 80% methanol at RT for 60 min followed by 6 x 10 min washes with 100% acetone. The specimens were stained with 2% OsO_4_in acetone at RT for 60 min followed by incubation in 1% UA in acetone + 10% methanol in the dark at RT for 60 min. After 4 washes in acetone for 15 min each they were infiltrated with increasing concentrations of Durcupan resin in acetone followed by incubation in pure Durcupan and polymerization at 60°C. For quality control the blocks were sectioned with a Leica LC6 ultramicrotome (Leica microsystems) and 300 nm sections were put on slot grids containing a Formvar film. Tilt series ranging from −63° to +63° were acquired with a F30 electron microscope (Thermo Fisher Scientific) and reconstructed with the IMOD software package^22^.

#### 2. Drosophila brain samples (fan-shaped body and accessory calyx)

While thin tissue samples (< 200-µm-thick) can be cryo-fixed, in many cases thick tissue specimens can not (e.g., large brain tissue) thus requiring various chemical fixation and staining protocols. *Drosophila* brain tissues from 5-day-old adult (Genome type: iso Canton S G1 x w1118 iso 5905) were prepared according to Progressive Lowering of Temperature and Low Temperature Staining (PLT-LTS) progressive heavy metal enhancement protocol described previously^23^. In details, after tissue dissection and pre-fixation, we osmicated tissue in 1% OsO_4_, then 1.5% K ferrocyanide, followed by a complete wash. Afterwards, a transfer to 1% thiocarbohydrazide for 15 min at 22°C, then a complete wash, followed by 2% osmium for 30 min at 22°C. After osmication, we stained in lead aspartate for 30 min at 55°C first, then 1 h at 22°C. Finally, the tissue was dehydrated in a Leica AFS freeze-substitution chamber: the temperature was dropped from 4°C to −25°C, and the concentration of acetone or ethanol was increased for 20 min in each of 10%, 30%, 50%, 70%, 80%, 90%, and 97%. The subsequent low temperature *en bloc* staining was performed in either 0.2% uranyl acetate in acetone, or 1% EPTA in 97% ethanol. Specimens were infiltrated and embedded in Durcupan.

### Preparation for FIB-SEM

After Durcupan re-embedding, a 3D X-Ray tomogram of the entire block was taken using an XRadia-510 Versa micro X-Ray system (Carl Zeiss X-ray Microscopy, Inc.). The 3D X-Ray tomograms allowed for robust Region of Interest (ROI) selection. Specifically, it identified cells with good morphology (for example a properly shaped cell, a cell in a desired stage of mitosis, a T-cell attacking a cancer cell, etc.), and also determined the proper orientation for the tissue samples. Once a potential ROI was identified, the sample block was then re-mounted to the top of a 1 mm copper post (using Durcupan) which was in contact with the metal-stained sample for better charge dissipation, as previously described^5^. Each sample was oriented on the copper post to allow the shortest distance along FIB milling direction. A small vertical sample post was trimmed to the ROI with a width of ∼100 µm and a depth of 60–80 µm in the direction of the ion beam for each sample. Sample trimming used an ultramicrotome (EM UC7, Leica Microsystems), was done in few iterations with X-Ray tomogram collected after each step. Once the desired ROI was trimmed, a thin layer of conductive material of 10-nm gold followed by 100-nm carbon was sputtered using a Gatan PECS 682 High-Resolution Ion Beam Coater. The coating parameters were 6 keV, 200 nA on both argon gas plasma sources, 10 rpm sample rotation with 45-degree tilt.

### FIB-SEM system advances

A Zeiss Gemini 500 and multiple Zeiss Merlin SEM systems were customized for this work to enable the two orders of magnitude improvement in imageable volume (from ∼500 µm^3^ to greater than 100,000 µm^3^) compared to our prior work^5^. Such improvement primarily comes from: 1) higher precision and stability of FIB milling control to extend reliable long-term acquisition to 4-nm voxels, and 2) enhanced SEM signal detection using secondary electrons to achieve faster imaging, hence reduce electron radiation energy density (from ∼400 to ∼80 keV/nm^3^) that further improves FIB milling control.

Specifically, to ensure precise and stable milling at 4-nm sampling, we integrated a more stable FIB column (Capella from Zeiss) and repositioned at 90 degrees to the SEM column (Fig. 1a). The Capella FIB column provided higher practical milling currents up to 30 nA compared to 7 nA of the Magnum FIB column (Thermo Fisher Scientific, previously FEI) used in our prior work^5^. We further tightened its FIB emission current control band from ± 100 pA to ± 50 pA to obtain a consistent milling beam profile. We also re-configured the milling closed-loop control (Fig. 10 in Xu et al., 2017^5^) to further reduce the milling variation between FIB reheat cycles thus enabling long-term acquisition at high electron radiation energy density. This FIB milling optimization sufficiently reduced milling artifacts hence allowed for the switching of SEM signal detection from backscattered electrons to secondary electrons for faster imaging (600 V vs. 0 V specimen bias in Fig. 12 of Xu et al., 2017^5^). The new SEM signal detection scheme lowered electron radiation energy density by 5x (from ∼400 to ∼80 keV/nm^3^), which in turn further mitigated FIB milling artifacts and instability. In addition, customized 100-µm-thick molybdenum apertures were laser machined to define more accurate and consistent FIB milling currents, and to achieve more than 2x longer lifetime; and the custom NI LabVIEW control software was re-designed: to include new configurations of FIB milling closed-loop control which improve milling variation; and to switch from the Zeiss RemCon serial port communication protocol to the Zeiss API which reduced the overhead by several seconds per imaging/milling cycle.

The optimization of SEM imaging conditions was then guided by the following principles. The isotropic resolution limits are convoluted by the waist size of the incoming primary electron beam and the scattering volume of the penetrating primary electrons. While spherical and chromatic aberrations of the electromagnetic lenses can be mitigated by smaller beam currents at modest numerical aperture to achieve the best beam focus and x-y resolution, lower primary beam landing energy generates fewer scattering events, resulting in smaller scattering volume which improves resolutions along all three axes. However, any further reduction below the optimal energy will result in a reduced scattering ratio of the heavy metal stain atoms over lighter background atoms of the sample resin, thus fading contrast. Through investigating a multi-dimensional operation space, we found that a beam current of 200 to 300 pA and a landing energy of 700 to 900 eV are optimal for isotropic 4-nm imaging.

Such positive synergy between faster SEM scanning and robust FIB milling has extended reliable imaging acquisition at 4-nm isotropic voxels from less than a week to months. Primarily limited by time, the maximum volume can be seamlessly extended.

### FIB-SEM imaging

As summarized in Table 1, all samples were imaged by a 0.14–0.3 nA electron beam with 0.7– 1.2 keV landing energy at 0.1–2.0 MHz. In most cases, both backscattered and secondary electron signals were collected by InLens detector to provide better signal-to-noise ratio. The time to acquire these volumes ranged from two to five weeks of uninterrupted imaging. Faster SEM scanning rates (5x–10x scanning speed improvement) were applied on *Drosophila* samples with stronger staining contrast while maintaining similar image quality, thus achieving maximum volume of 175,000 µm^3^ within three-week imaging.

For all samples, the x-y pixel size was set at 4 nm. A subsequently applied focused Ga+ beam of 15 nA at 30 keV strafed across the top surface and ablated away 4 nm of the surface. The newly exposed surface was then imaged again. The ablation – imaging cycle continued for two to five weeks of uninterrupted imaging to complete FIB-SEM imaging one sample. The sequence of acquired images formed a raw imaged volume, followed by post processing of image registration and alignment using a Fiji plugin based on Scale Invariant Feature Transform (SIFT) algorithm^24^ to form isotropic 4 nm voxels. The final aligned stack consisted of an isotropic volume, containing multiple complete cells, which can be viewed in any arbitrary orientations.

### Evaluation of FIB-SEM resolution

In order to evaluate the actual resolution in the acquired FIB-SEM datasets we analyzed the transitions of the edges of the ribosomes abundant within the cell volumes. Ribosomes are macromolecular machines consisting of RNA and associated proteins. Their small size (20–30 nm), abundance in the cytoplasm of living cells, and the fact that they are stained with high EM contrast makes them good candidates for resolution evaluation.

First, we selected volumes of approximately 1000–2000 x 1000 x 1000 pixels in cultured cell datasets. These volumes were selected in the middle of datasets in areas with large number of ribosomes. We then calculated histograms of grey level values in these volumes and subtracted ∼0.8 * histogram max to bring the average cytoplasmic material signal level close to 0. We then used the Laplacian of Gaussian (LoG) algorithm (part of Python scikit-image package^25^) to select blobs. We set the limit of standard deviation for LoG between 1 and 2.5 pixels (all these datasets have 4-nm voxels) to ensure that selected blobs were predominantly ribosomes. Filters were used to exclude the following blobs: average value below zero (this meant LoG algorithm failed), edge value above 0.4 times of max value or 37%–63% transition longer than 15nm (the latter two happen where there is another blob or another feature with high signal level in close proximity—making subsequent edge transition analysis inaccurate), amplitudes below certain threshold, so that at least approximately 3000 blobs per dataset were remained for analysis. The edge transitions in all three directions were analyzed for these remaining blobs, in particular we determined 37%–63% transitions (the value used by Zeiss in their resolution estimation) and 20%–80% transitions (close to 1 sigma value).

The summaries of the 37%–63% transitions and 20%–80% transitions for the 3D ribosome blobs selected in cultured cell datasets are plotted in Fig. 1b. The histograms of the 37%–63% transitions for these datasets are shown in Extended Data Fig. 3. The examples of 3D blobs for each dataset are presented in Extended Data Fig. 4–10. For each dataset the cross-sections and transition analysis for 3 brightest and 3 dimmest ribosomes are shown.

### Accurate determination of FIB-SEM milling rate

The position of a sample block face was measured by the SEM beam and the FIB beam independently. An in-line auto focus routine^6^ kept the SEM images in focus throughout the entire image acquisition. While the working distance of the SEM beam detected the block face position by measuring its distance from the SEM objective lens, the FIB milling position under closed-loop control indicated the block face position directly from the orthogonal angle. As the sample block face being imaged then milled away layer-by-layer, the SEM working distance and the FIB beam position tracked the block face location accordingly. Therefore, we estimated the total thickness milled away by FIB based on the changes of either SEM working distance or FIB milling position from the beginning to the end of the FIB-SEM acquisition. The average milling rate of z-step was then calculated as the total milling depth divided by the number of image frames. Here we report the average z-scaling factors derived from both measurement approaches in Extended Data Table 1, which can be used to scale the dataset to generate real isotropic voxels. For example, HeLa Cell 2017-06-21 dataset has a z-scaling factor of 1.31 which means the actual z-step is averaged at 5.2 nm instead of 4.0 nm.

### Image emulation to visualize the value of finer resolution

To emulate 8-nm images, we first applied a 6-nm Gaussian blur filter and then added Gaussian noise with a standard deviation of 22 to 4-nm images. The higher shot noise at 8-nm sampling was expected due to the 10x lower electron dose of ∼12 e/nm^3^, compared to that of ∼120 e/nm^3^ at 4-nm sampling. The 6-nm Gaussian blur and added noise (Fig. 1d, g, and j) were underestimated compared to that of the real 8-nm images (Fig. 1e, h, and k) collected on the same type of specimens. The 8-nm emulated images with comparisons to 4-nm images and real 8-nm images are shown in Fig. 1c–k and Supplementary Video 1.

## Data availability

All raw and segmented datasets in this work are available on OpenOrganelle website (https://openorganelle.janelia.org).

## Code availability

FIB-SEM image acquisition LabVIEW code used in this work is available from https://github.com/cshanxu/Enhanced_FIB-SEM

Python code for resolution characterizations using ribosomes is available from https://github.com/gleb-shtengel/FIB-SEM_resolution_evaluation

## Acknowledgements

We thank K. Hayworth and W. Qiu at Howard Hughes Medical Institute (HHMI) Janelia Research Campus (JRC) for invaluable discussions and data collection support. We gratefully acknowledge P. Rivlin, S. Plaza, and I. Meinertzhagen for JRC EM shared resource and FlyEM project team support on staining protocols development. We thank the electron microscopy facility of MPI-CBG and of the CMCB Technology Platform at TU Dresden for their services. We also thank Y. Wu from Pietro De Camilli’s laboratory at Yale for advice.

## Author Contributions

C.S.X. and H.F.H. supervised the project; C.S.X., S.P., G.S., and H.F.H. wrote the manuscript with input from all co-authors; C.S.X. developed enhanced FIB-SEM platform for large volume high resolution imaging, and optimized imaging conditions; C.S.X., S.P., and G.S. conducted FIB-SEM experiments; C.S.X. and S.P. performed image post processing; S.P., G.S., A.M., A.T.R., H.K.H., S.B.v.E., Z.L., H.A.P., N.I., J.C., A.V.W., and M.F. prepared samples; G.S., A.M., A.T.R., and S.T. analyzed data; D.B. prepared and uploaded data to OpenOrganelle website; S.B.v.E., T.C.W., R.V.F., J.L.-S., I.M., and M.S. proposed biological questions and provided samples.

---

**Extended Data Fig. 1.**
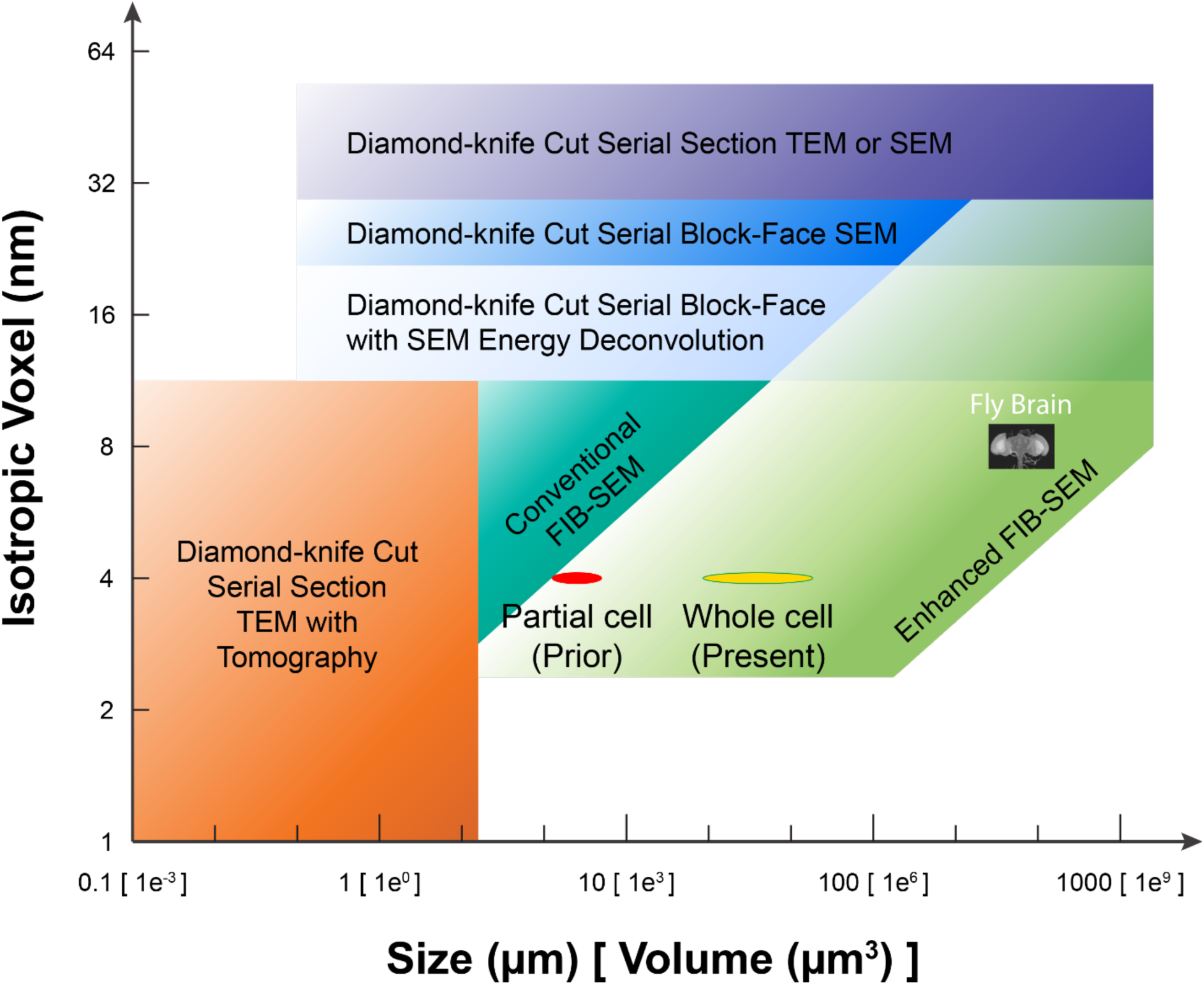
Isotropic voxel (representing the minimal voxel size dictated by the worst-case axial resolution) vs. volume for comparing different volume EM methods. The light green space represents the Resolution-Volume regime accessible with enhanced FIB-SEM technology through long term imaging. The present work of whole cell volumes at 4-nm isotropic voxels matching the resolutions shown in Fig. 1b is colored in yellow, compared to the prior work of smaller volumes colored in red. Adopted from Xu et al., 2017^5^ with modifications.

**Extended Data Fig. 2.**
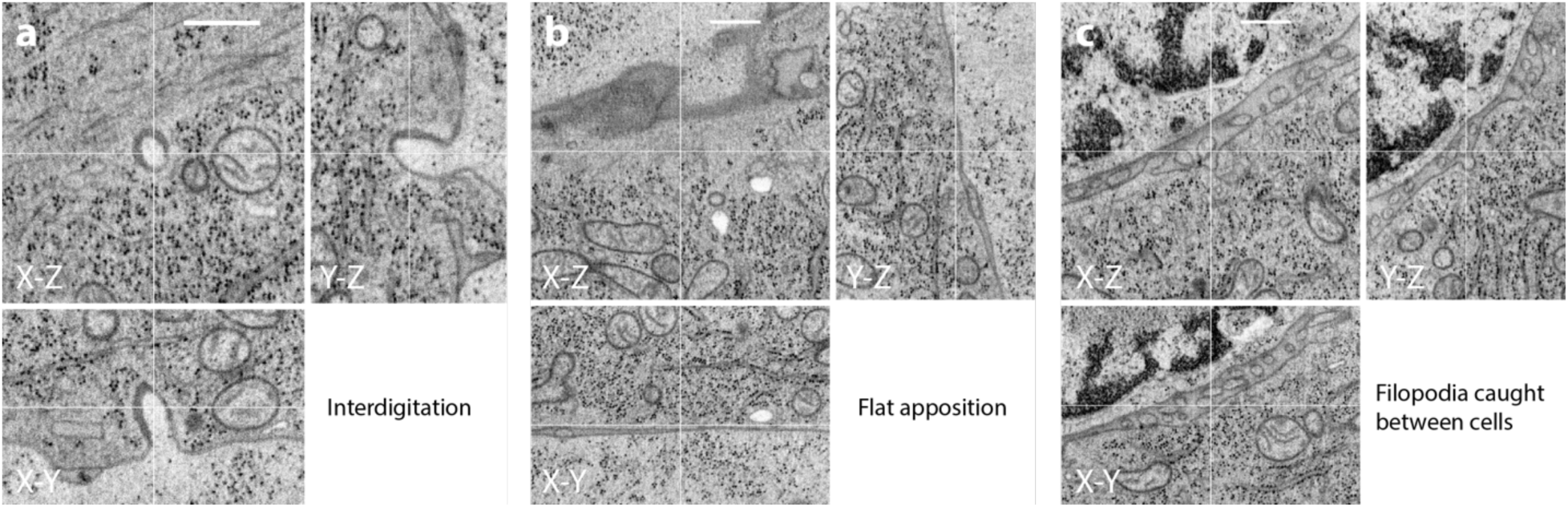
Murine CTL engaging an ovarian cancer cell. Zooms on regions showing different immunological synapse topology features. **a,** Interdigitation, **b,** Flat apposition, and **c,** Filopodia caught between cells. Scale bars are 0.5 μm.

**Extended Data Fig. 3.**
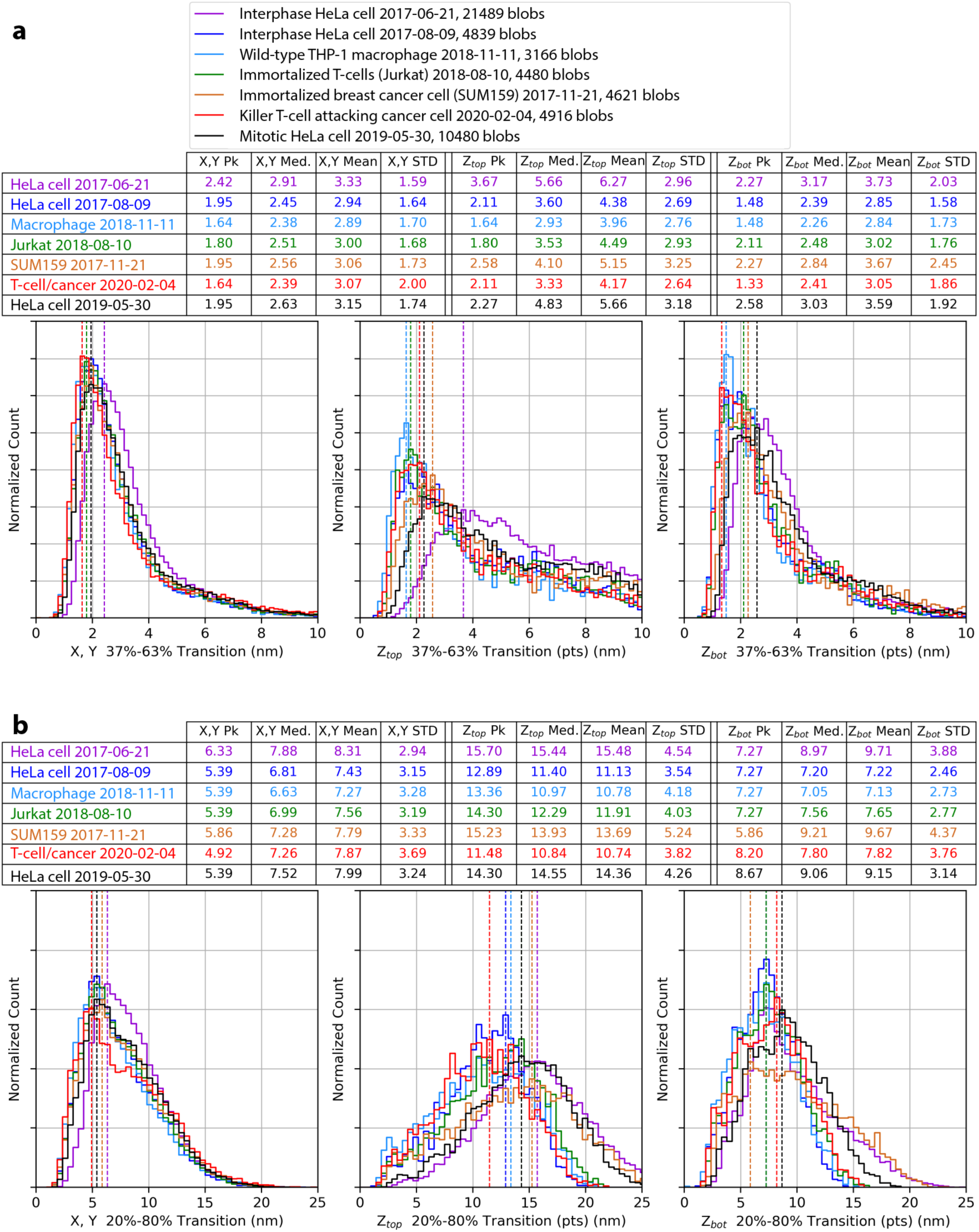
Edge transition distributions determined from ribosomes in cultured cells datasets. Distributions of 37%–63% (**a**) and 20%–80% (**b**) transition distances in X-, Y-(left), Z_top_-(center), and Z_bot_-(right) directions.

**Extended Data Fig. 4.**
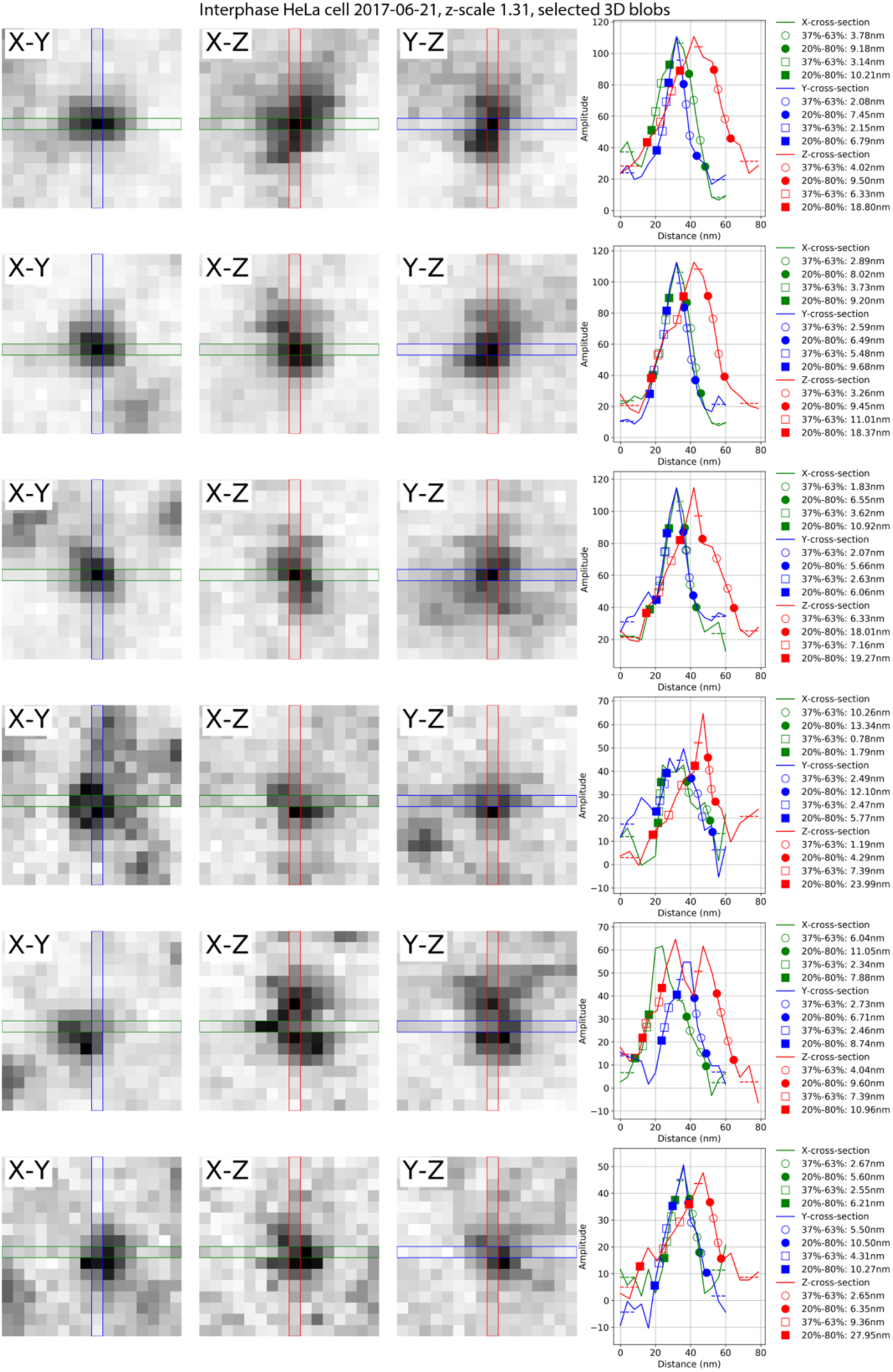
Cross-sections of the example of ribosomes from the dataset Interphase HeLa Cell 2017-06-21 and the profiles with the transition analysis. The top three rows are the brightest ribosomes and the bottom three rows are the dimmest ribosomes.

**Extended Data Fig. 5.**
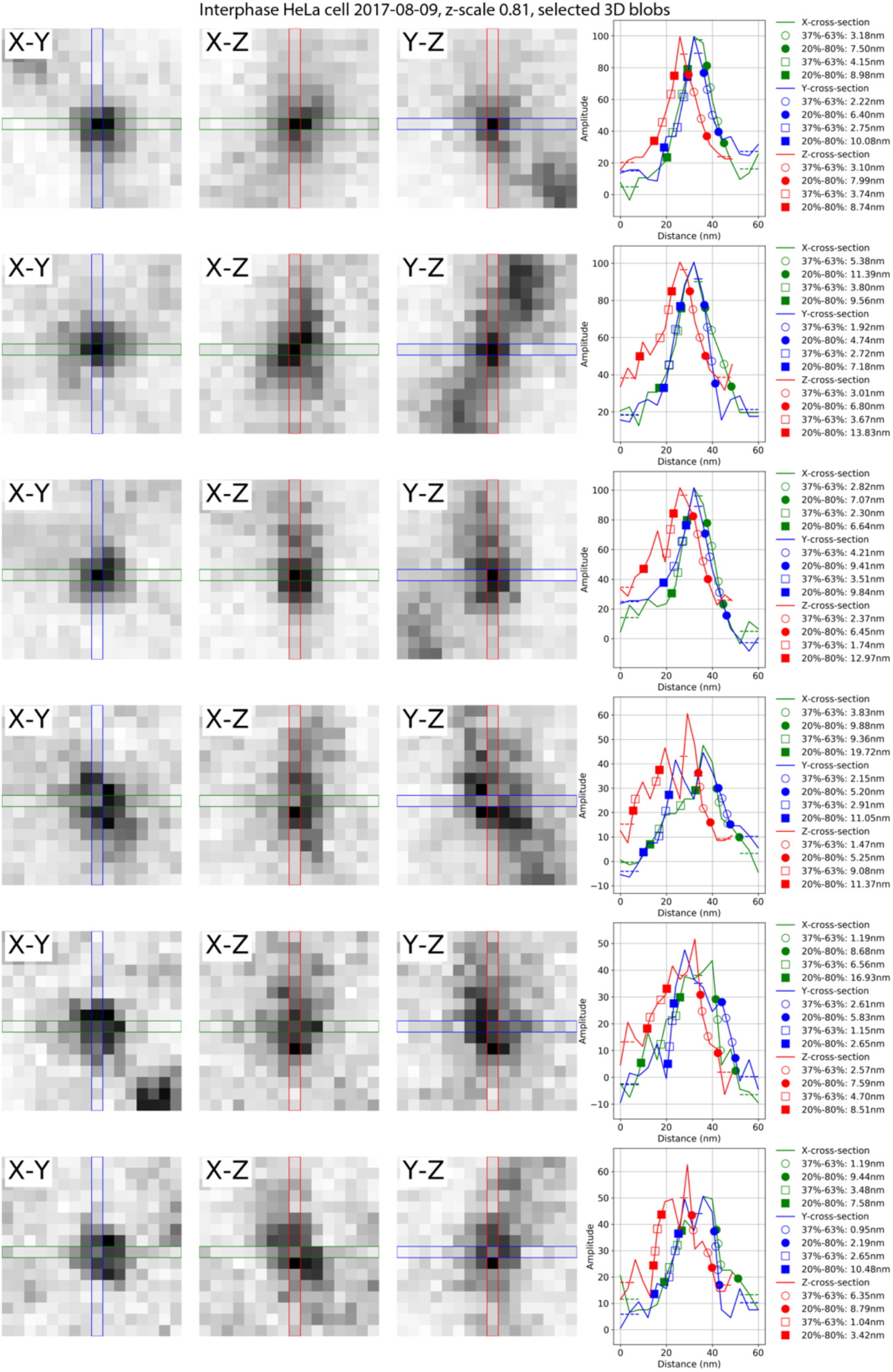
Cross-sections of the example of ribosomes from the dataset Interphase HeLa Cell 2017-08-09 and the profiles with the transition analysis. The top three rows are the brightest ribosomes and the bottom three rows are the dimmest ribosomes.

**Extended Data Fig. 6.**
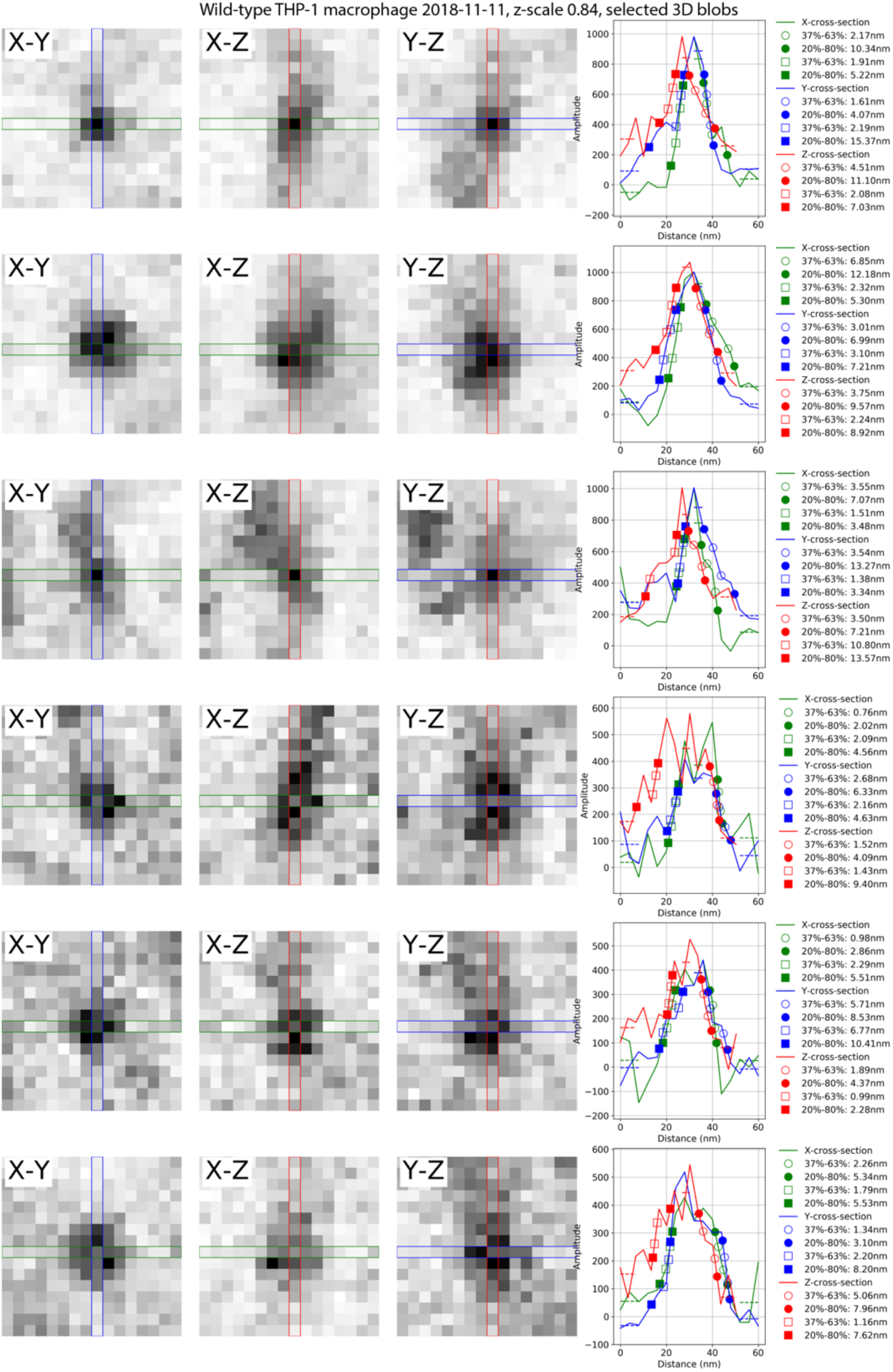
Cross-sections of the example of ribosomes from the dataset Wild-type THP-1 Macrophage 2018-11-11 and the profiles with the transition analysis. The top three rows are the brightest ribosomes and the bottom three rows are the dimmest ribosomes.

**Extended Data Fig. 7.**
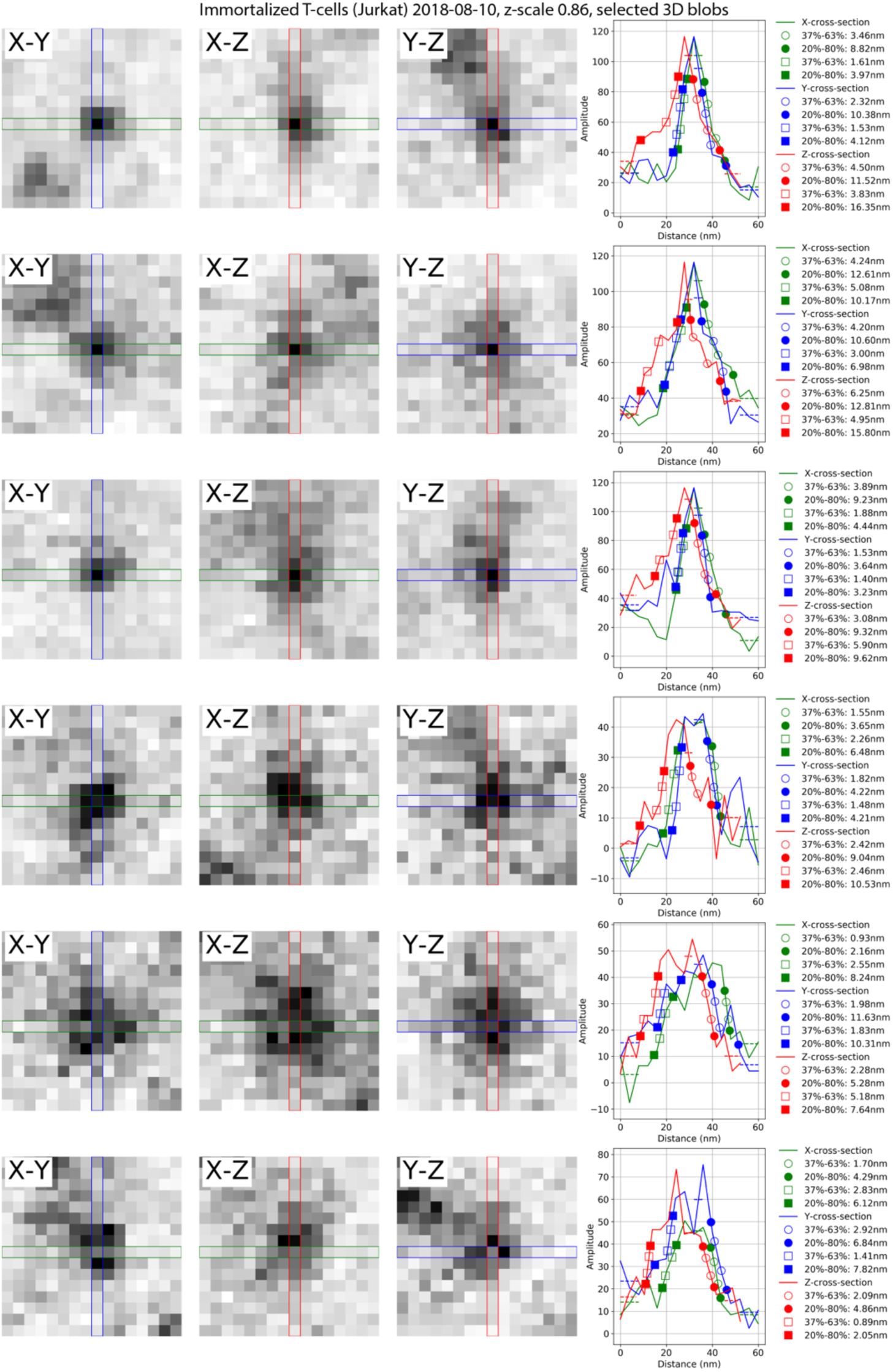
Cross-sections of the example of ribosomes from the dataset Immortalized T-cells (Jurkat) 2018-08-10 and the profiles with the transition analysis. The top three rows are the brightest ribosomes and the bottom three rows are the dimmest ribosomes.

**Extended Data Fig. 8.**
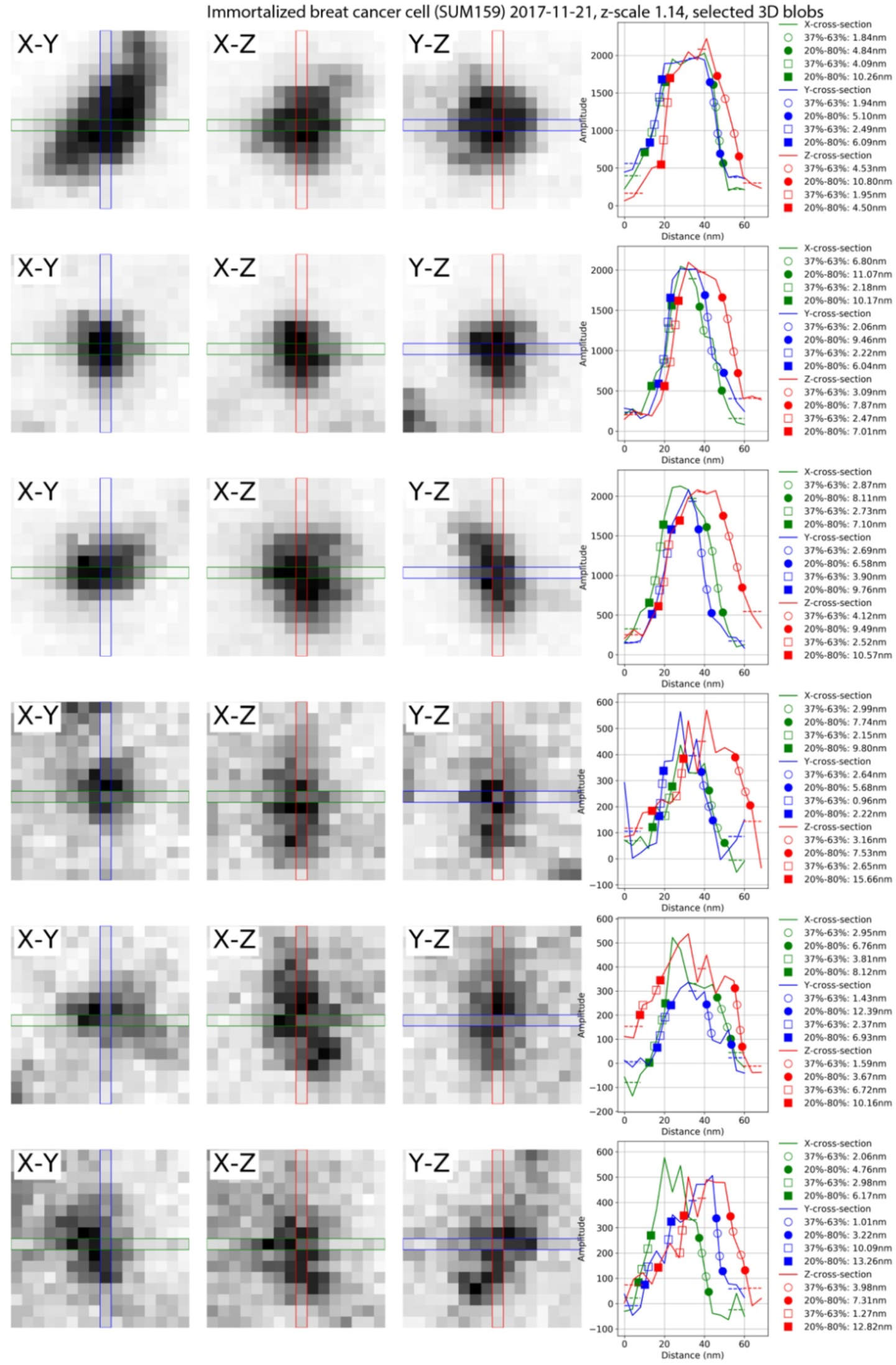
Cross-sections of the example of ribosomes from the dataset Immortalized breast cancer cell (SUM159) 2017-11-21 and the profiles with the transition analysis. The top three rows are the brightest ribosomes and the bottom three rows are the dimmest ribosomes.

**Extended Data Fig. 9.**
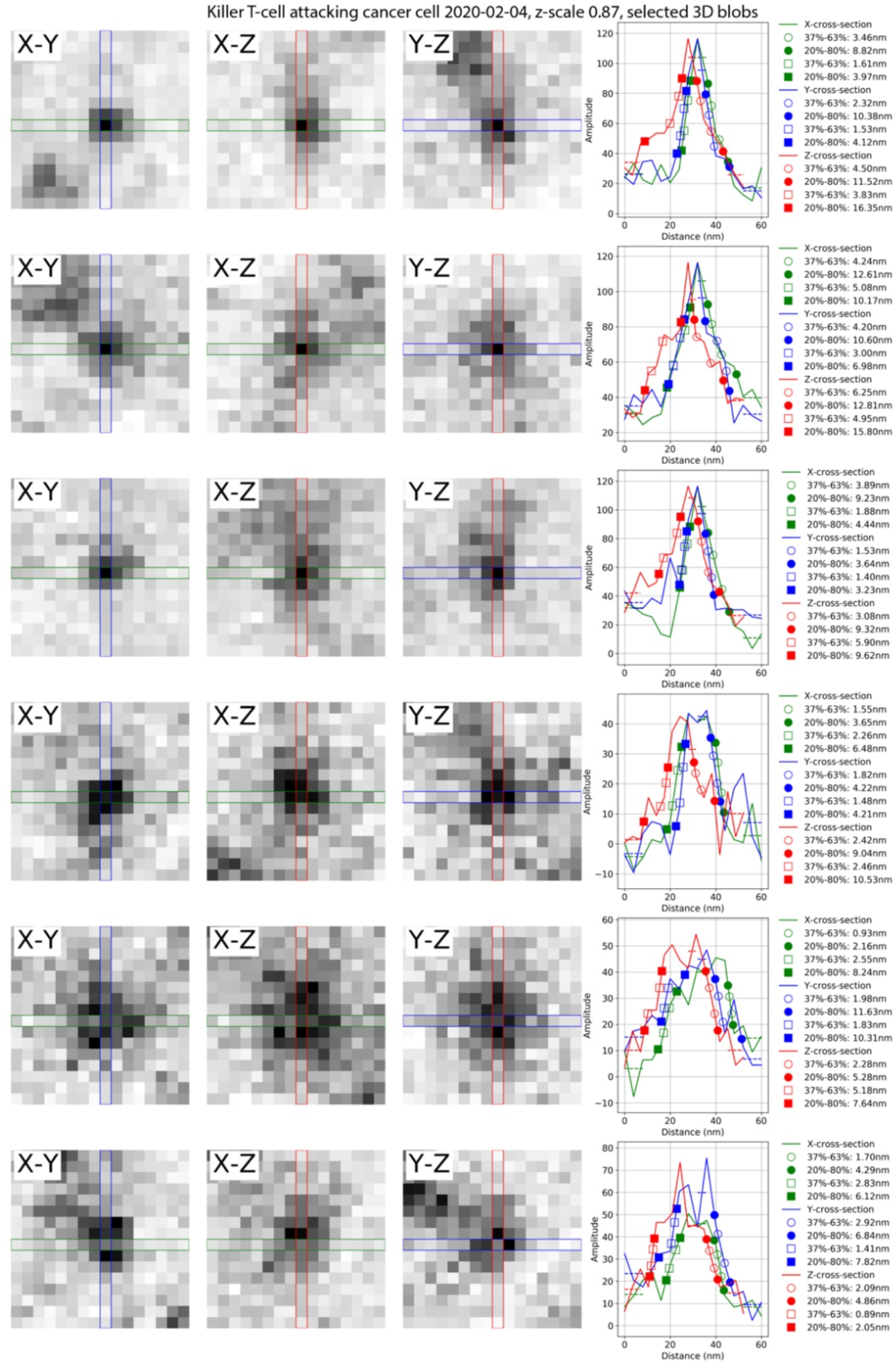
Cross-sections of the example of ribosomes from the dataset Killer T-cell attacking cancer cell 2020-02-04 on Cancer Cell and the profiles with the transition analysis. The top three rows are the brightest ribosomes and the bottom three rows are the dimmest ribosomes.

**Extended Data Fig. 10.**
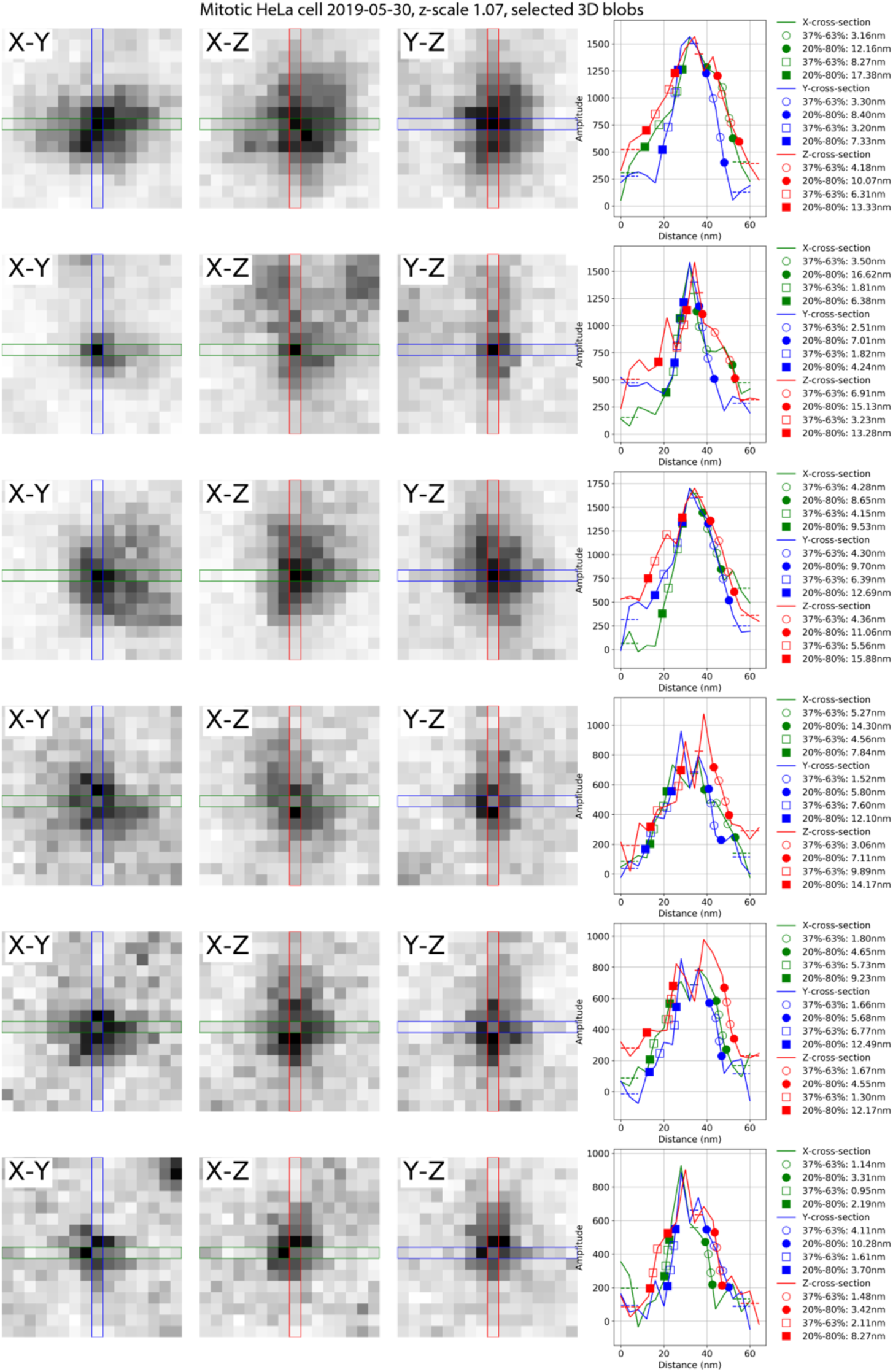
Cross-sections of the example of ribosomes from the dataset Mitotic HeLa cell 2019-05-30 and the profiles with the transition analysis. The top three rows are the brightest ribosomes and the bottom three rows are the dimmest ribosomes.

**Extended Data Table 1.**
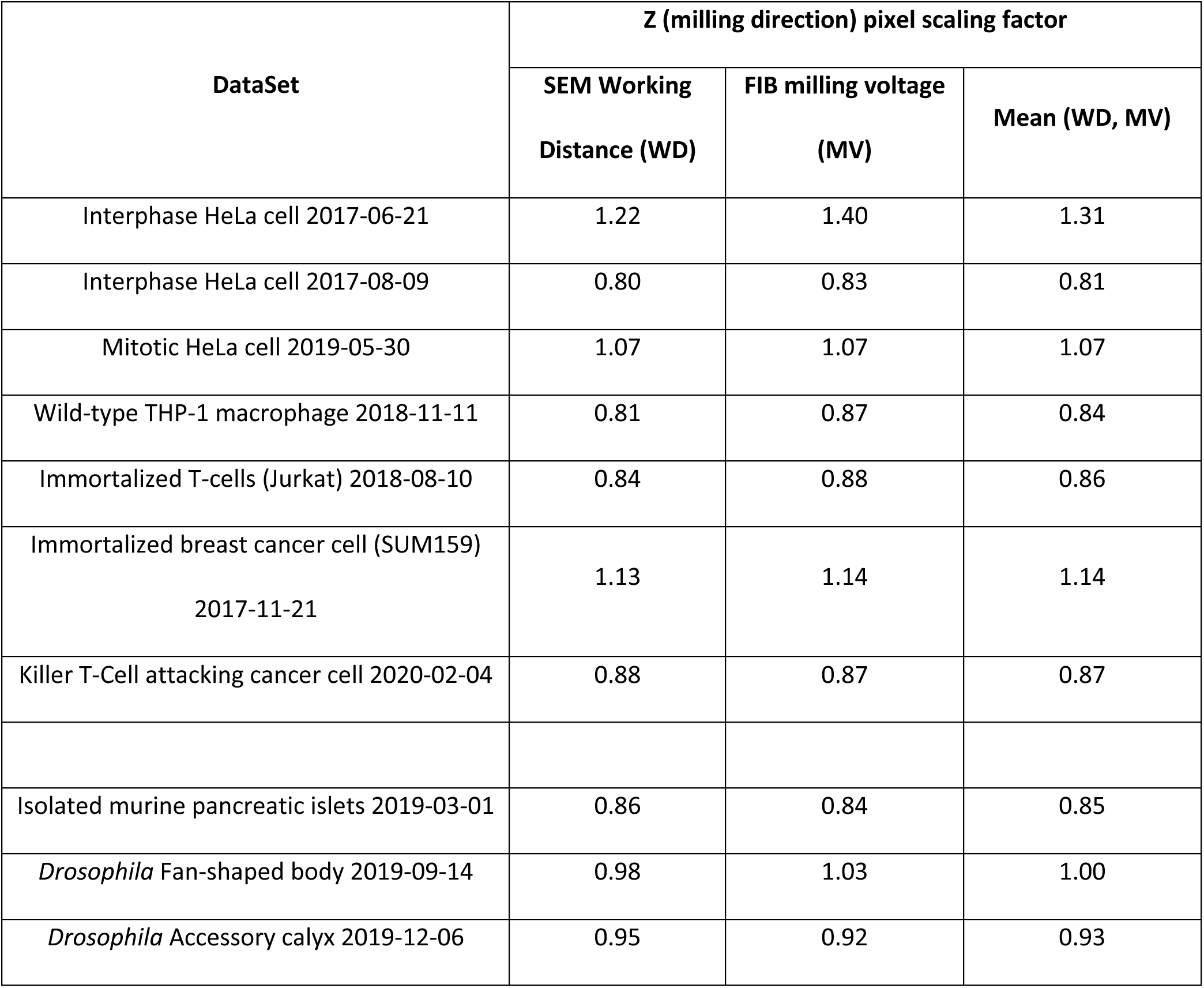
. Z scaling factors estimated using SEM working distance and FIB milling voltage.

