## Supplementary Information for "An open-access volume electron microscopy atlas of whole cells and tissues"

### Supplementary Methods

#### Cultured cell sample preparation

##### 1. Coverslip preparation

Sapphire coverslips (3 mm diameter, 0.05 mm thickness, Nanjing Co-Energy Optical Crystal Co., Ltd) were cleaned in basic piranha solution (5:1:1 solution of water : ammonia hydroxide : hydrogen peroxide (30%)) for a minimum of 1 hour. A thin (~100 nm) coat of gold was sputtered (sputter coater Desk II, Denton Vacuum) on to a 0.5 mm wide border region of clean coverslips, differentiating the surfaces such that the sides with cells plated on them (opposite of the gold coated sides) could be identified at all steps in the process.

##### 2. Freezing

Cultured cell samples were cryo-fixed using a Wohlwend Compact 2 high-pressure freezer (Engineering Office M. Wohlwend GmbH, Bifig 14, 9466 Sennwald, Switzerland). Immediately prior to freezing, live cells are inspected to ensure cell morphology and viability. After quality assurance the cells are transferred to a water jacketed $CO_2$ incubator (ThermoFisher, Midi 40) kept at 37° C, 5% $CO_2$, and 100% humidity while awaiting freezing. Each coverslip is removed from the incubator immediately prior to the freezing procedure. The freezing procedure consists of eight steps[1]:

a. Coat the sides of the aluminum platelets that will contact the sapphire (coverslip) or cell media with hexadecene. We use Technotrade International, Alu Platelet 479 (Cavity 0.3 mm with one side ground) and 389 (Specimen Carrier Cavity 0.025 / 0.275 mm) for freezing on a 3-mm diameter 50-µm-thick sapphire coverslip. Place

479 disks ground (flat) side down, and 389 disks 0.025mm side down to coat hexadecene.

b. Blot the hexadecene from the platelets with filter paper (Whatman® qualitative filter paper, Grade 1)
c. Prepare the HPF holder by placing the flat platelet (Technotrade International, Alu Platelet 479) into it with ground (flat) side facing up.
d. Remove a coverslip from the incubator
e. Replace the media on the coverslip with a 25% w/v Bovine Serum Albumins (BSA, Sigma, B4287) media mixture by dipping the coverslip (cell side down) into three independent 20 µL reservoirs of the BSA mixture
f. Place the coverslip (cell side down) with the replaced media onto the 25 µm well of aluminum platelet (Technotrade International, Al Platelet 389 (0.025/0.275))
g. Blot this “sandwich” with filter paper to remove the excess media and then place it into the holder such that the back side of coverslip sits on the ground (flat) side of Alu Platelet 479.

   Please note, as a result from steps f and g, the back side of coverslip sits on the ground (flat) side of Alu Platelet 479, and cell side of coverslip faces up onto the 25 µm well of Al Platelet 389.
h. Freezing is completed by following the manufacturer’s instructions and the frozen sandwich is stored under liquid nitrogen in a custom-made holder manufactured out of brass or aluminum

The BSA mixture is used as a cell cryo-protectant to prevent ice crystal formation in the extra cellular space[2]. Once the sample is frozen it can be stored indefinitely in liquid nitrogen for subsequent freeze-substitution and resin embedding.

#### 3. Freeze-substitution and resin embedding

The coverslips with high-pressure frozen samples are prepared for room temperature electron microscopy by freeze substitution and heavy metal staining followed by embedding in resin.

##### *a. Freeze-substitution*

Freeze-substitution (FS) was performed with a protocol adapted from[3]. Briefly, coverslips were transferred to cryotubes containing FS media (2% $OsO_4$, 0.1% Uranyl Acetate (UA), and 3% water in acetone) under liquid nitrogen and the following FS schedule was executed using automated FS machine (AFS2, Leica Microsystems):

i) 140°C to -90°C 2h
ii) -90°C to -90°C 24h
iii) -90°C to 0°C 12 h
iv) 0 to 22°C 1h
v) 22°C 1 h

##### *b. Resin Embedding*

Resin embedding was performed immediately after FS. Samples were removed from the AFS2 machine, washed 3 times in anhydrous acetone for a total of 10 min and embedded in Eponate 12 with BDMA (Ted Pella, kit 18012) with the following protocol:

i) Acetone/Eponate 12 2:1 1h
ii) Acetone/Eponate 12 1:1 1 h

iii) Acetone/Eponate 12 1:2 1 h

iv) Eponate 12 2 h

v) Eponate 12 2 h

vi) Eponate 12 2 h

Coverslips were placed in the slots of a flat embedding silicone mold, cells side up, and immersed in Eponate 12 which was polymerized for 48 hours at 65°C.

##### *c. Re-Embedding*

Following EPON embedding, the epoxy is removed from all coverslip surfaces not containing cells using a razor blade (RICA Surgical Products, Wec-Prep Blades, 450205). Then the coverslip is separated from the resin block containing the cells by sequential immersion in liquid nitrogen and hot water. In a process similar to pothole formation, hot water gets into the cracks between the resin and the coverslip and then expands when it freezes in the liquid nitrogen. Moreover, the sapphire coverslip and polymer resin have very different thermal expansion coefficients. Both mechanisms combine to ensure easy coverslip separation. EPON is chosen as the initial embedding resin due to less embedding artifacts. However, EPON generates streaking artifact during FIB-SEM imaging[4]. A thin layer (5–10 µm) of Durcupan on the specimen surface facing the FIB beam can effectively mitigate such streaks during FIB-SEM imaging. Therefore, once the coverslip is removed, the exposed surface is immediately re-embedded in Durcupan ACM resin (Sigma Aldrich, set 44610) before trimming to the region of interest.

#### Supplementary Videos

Supplementary Video 1. The FIB-SEM dataset of *Drosophila* brain fan-shaped body imaged at 4-nm voxels with 100,000 $\mu m^3$, demonstrates ~200x larger volume compared to that of a differently stained *Drosophila* brain protocerebral bridge sample with volume of ~500 $\mu m^3$ reported in prior work[4]. Finer ultrastructures (e.g., ER-Mito contact and microtubule hollow core) revealed at 4-nm voxels are not distinguishable at the emulated 8-nm voxels. The emulated 8-nm condition is underestimated compared to that of the real 8-nm images collected on the same type of specimens shown in Fig. 1i–k.

Supplementary Video 2. Fine 3D structural details of immunological synapse (T-cell "cupping" cancer cell, polarized centrosome, filopodia trapped between two cells, and lytic granules), and its surrounding in CTL attaching target cell.

https://doi.org/10.7554/eLife.25916
